# Utilizing Transcranial Direct Current Stimulation to Enhance Laparoscopic Technical Skills Training: A Randomized Controlled Trial

**DOI:** 10.1101/455329

**Authors:** Morgan L. Cox, Zhi-De Deng, Hannah Palmer, Amanda Watts, Lysianne Beynel, Jonathan R. Young, Sarah H. Lisanby, John Migaly, Lawrence G. Appelbaum

## Abstract

This study aimed to test the efficacy of transcranial direct current stimulation (tDCS) during laparoscopic skill training to determine if it has the capacity to accelerate technical skill acquisition. tDCS is a non-invasive brain stimulation technique that delivers constant, low electrical current resulting in changes to cortical excitability and prior work suggests it may enhance motor learning. We evaluate for the first time the potential of tDCS, coupled with motor skill training, to accelerate the development of laparoscopic technical skills. In this pre-registered, double-blinded and sham-controlled study, 60 healthy subjects were randomized into sham or active tDCS in either bilateral primary motor cortex (bM1) or supplementary motor area (SMA) electrode configurations. All subjects practiced the Fundamental of Laparoscopic Surgery Peg Transfer Task during a pre-test, six 20-minute training sessions, and a post-test. The primary outcome was change in laparoscopic skill performance over time, quantified by improvement in performance according to a seconds-per-object calculated score accounting for errors. Sixty participants were randomized equally into the three training cohorts (active bM1, active SMA, sham). The active groups had significantly greater improvement in performance from pre-test to post-test compared to the sham groups (108 vs 76 seconds, *p* = 0.018). Both bM1 and SMA active cohorts had significantly greater improvement in learning (*p* < 0.01), achieving the same skill level in 4 sessions compared to the 6 sessions required of the sham cohort. The SMA cohort had more variability in performance compared to the bM1 and control cohorts. Laparoscopic skill training with active, bM1 or SMA, tDCS exhibited significantly greater learning relative to training with sham tDCS. The potential for tDCS to enhance the training of surgical skills merits further investigation to determine if these preliminary results may be replicated.

## I. INTRODUCTION

The field of surgery has continued to evolve over time with technological advances, work hour restrictions, and emphasis on patient safety [1–3]. This evolution has driven surgical residencies to focus on simulation and competency-based training to target technical skill development in a safe, controlled environment prior to patient interaction [4–8]. The importance of this training paradigm has been exemplified by the American Board of Surgery requiring that all general surgery residents pass the one-time, simulation-based certifications in Fundamentals of Laparoscopic Surgery (FLS) and Fundamentals of Endoscopic Surgery (FESTM) [9, 10]. The development of these basic technical skills requires the removal of residents from the bedside in order to achieve repetition of training within the simulation lab [11–14]. Therefore, it is imperative that new technologies and training techniques are developed symbiotically to expedite the time-consuming development of basic technical skills in the simulation lab.

Transcranial direct current stimulation (tDCS) is a non-invasive brain stimulation technique that delivers constant, low electrical current via saline-soaked electrodes on the scalp to modify neuronal excitability in the underlying cortex [15–17]. Anodal current has been shown to increase cortical excitability and facilitate spontaneous neuronal firing [18]. Numerous studies suggest that tDCS to motor cortex, in some cases coupled with simultaneous motor skill training, may enhance motor learning in healthy individuals [19–23] as well as faster motor-system recovery following stroke in clinical populations [24, 25]. While a recent consensus paper on tDCS and motor learning described promising results, it also highlighted a pressing need for prospective, preregistered, hypothesis-based studies to improve the transparency and replication of this approach so that the potential of these interventions may be rigorously established [26]. The ease with which tDCS can be implemented and tolerated, paired with the potential for lasting changes in cortical excitability and associated motor learning, gives this approach the powerful potential to accelerate the development of surgical technical skills [27, 28]. Additionally, the safety profile of tDCS has been well described with parameters feasible for implementation within a surgical skills training lab [29–31].

Only one prior study has been published on the application of tDCS within surgery [32]. In that study, Ciechanski and colleagues applied anodal tDCS to 22 medical students performing a neurosurgical tumor resection task resulting in improved efficiency and effectiveness of performance. While this is the first evidence that tDCS can positively affect surgical skill training, the study was conducted within a surgical subspecialty and targeted a more advanced task. The application of tDCS within general surgery training, targeting fundamental and generalizable skills necessary to pass required technical skill assessments, has yet to be investigated. Additionally, given that tDCS has not previously been applied in a bimanual surgical skill task, the optimal electrode configuration for greatest performance enhancement remains unknown.

Therefore, the current study aimed to test the efficacy of tDCS during laparoscopic skill training to determine if it has the capacity to accelerate technical skill acquisition. We elected tDCS parameters to be in the range of those previously reported to affect motor skill training. Since the optimal electrode configuration is not known, we evaluated two separate electrode configurations, each targeting the motor system (bilateral primary motor area and supplemental motor area). We hypothesized active tDCS would lead to faster and larger learning gains relative to sham tDCS while performing the FLS Peg Transfer Task, and we evaluated the exploratory hypothesis that the two active placements would differ in their effects. We selected the SMA target due to its functional role in motor planning and execution, as well as bimanual coordination; prior tDCS studies have shown that stimulation of the SMA can increase performance on simple and complex motor tasks [33].

## II. METHODS

### A. Participants

This double-blinded, randomized, and sham-controlled study was approved by the Duke Institutional Review Board and pre-registered at ClinicalTrials.gov (NCT03083483).

Healthy adults with the ability to provide consent and follow directions were included in the study. Exclusion criteria were specific to tDCS including brain tumor, psychotropic medications, seizure disorder, substance abuse, and metal implants in the head. Females had to provide a negative urine pregnancy test at the initiation of the study. Subjects meeting inclusion criteria on a screening survey provided written consent and were enrolled into the study. Subjects were provided $20 per hour compensation for their time.

### B. Randomization

Prior to the start of the study, simple randomization was used to assign a tDCS code and electrode configuration to each participant. Each code was programmed as either active or sham stimulation by the device manufacturer, and this information remained blinded to the study team and subjects until after statistical analysis.

Electrode configurations targeted either the bilateral primary motor cortex (bM1) or the supplementary motor area (SMA) and balanced randomization was performed to assign subjects equally to active bM1, active SMA, and sham cohorts (half with bM1 and half with SMA configurations).

### C. tDCS Specifications

The Soterix Medical 1 × 1 tDCS Device (Soterix Medical Inc., New York, NY) was used. Active stimulation consisted of a 30-second, gradual ramping up of electrical current to a maximum sustained level of 2 mA for the duration of the 20-minute training session, followed by a 30- second down ramp to 0 mA current. Sham tDCS consisted of the 30-second on ramp, followed immediately by the off ramp to create an initial cutaneous sensation of stimulation without the sustained current delivery [34]. The device provided another 30-seconds of stimulation at the end of the 20-minute training session.

Electrodes were placed on each subject utilizing the 10–20 electroencephalography electrode system, with the two, 3 cm × 5 cm, saline-soaked sponges held in place using two Soterix Medical Elastic Fastener “Blue” bands (Soterix Medical Inc., New York, NY). For the bM1 configuration, the cathodal electrode was placed over C4 while the anodal electrode was placed over C3. SMA configuration placed the cathodal electrode over Fpz and the anodal electrode 15% of the nasion–inion distance anterior to vertex, with the long edge of the sponge pads oriented left–right. Electrode configurations and underlying cortical electric field illustrations are shown in Figure 1. Electric field simulation was performed using SimNIBS 2.1.1. [35].

**Figure 1:**
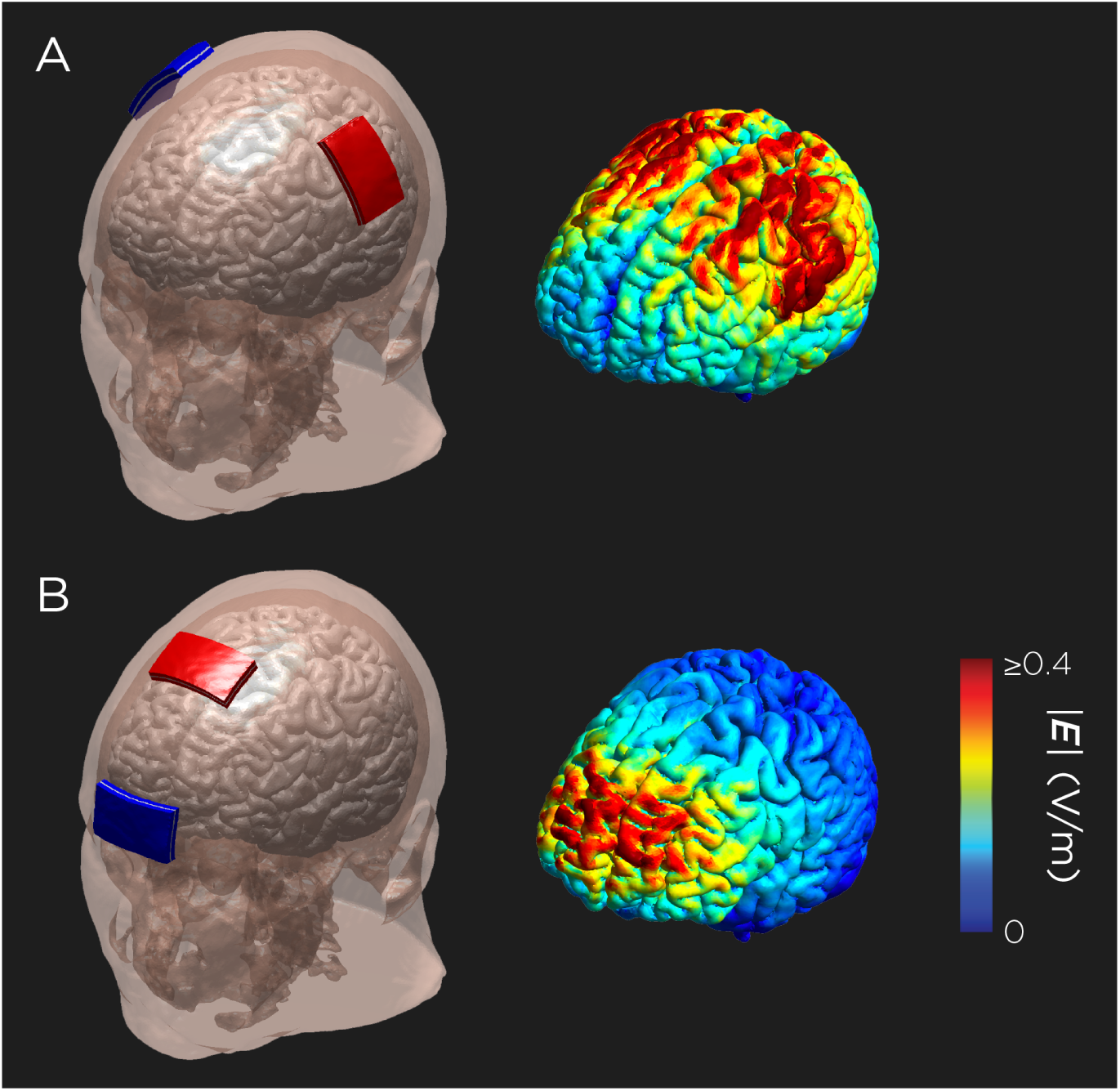
tDCS electrode (red = anode, blue = cathode) configurations and underlying electric field distribution in the brain for the A) bilateral primary motor cortex montage (bM1) and B) the supplementary motor area (SMA) montage.

Electrode conductivity was measured continuously throughout the session, and instances of increased contact impedance were noted and addressed by adding more saline to the sponges or increasing the amount of pressure applied by the elastic bands. If necessary, the session was stopped and restarted for repositioning of the electrodes. The occurrence and duration of any critical connectivity events were documented. No subject had to be excluded due to an inability to provide adequate tDCS.

### D. Manual Skills Training

Each subject viewed the same 4-minute online instructional video on how to perform the FLS Task 1 Peg Transfer (https://www.youtube.com/watch?v=gAQPXHWqdXQ). This was followed by approximately 10-minutes of acclimation to the FLS box trainer (Limbs & Things Ltd., Savannah, GA) with time allowing for one repetition of the Peg Transfer Task, consisting of moving all 6 triangle objects to the right and back to the left, while receiving verbal feedback from a study team member to point out any violations of the procedure.

Laparoscopic surgical skill training consisted of six separate 20-minute sessions where the subject repeatedly performed the FLS Peg Transfer Task. Training sessions were completed during three study visits within a 7-day window with two sessions completed each visit and a maximum five-minute break in between. Additionally, a pre-test and a post-test consisting of one repetition of the task were administered prior to and upon completion of the six training sessions. In the three instances where a pre or post-tests was not completed, the first repetition or last repetition was timed and included for pre-post analysis.

### E. Outcomes and Data Acquisition

Study data were collected and managed using Research Electronic Data Capture (REDCap) electronic data capture tools hosted at Duke University [36].

Subjects completed a questionnaire consisting of baseline characteristics, prior surgical skill experience, and personal behaviors which was repeated at the beginning of each visit. Baseline characteristics included age, sex, ethnicity, race, and handedness. Prior experience with either open or laparoscopic surgery was obtained. Personal behaviors included self-identification as an athlete, musician, video gamer, or artist in addition to servings of caffeine consumed and hours of sleep obtained the day of skills training.

Videos of each study visit were recorded using the Argus Science Frame Grabber (Argus Science, LLC, Tyngsborough, MA). This system digitizes the analog signal directly from the camera located within the FLS box trainer. Each video was reviewed and scored at a later date by a single study team member (MLC) with surgical experience to ensure internal consistency. Errors were defined as 1) drops of the object outside the field of view and 2) imprper transfers (transfer on a drop, transfer with the wrong hand, resting the object during transfer). These errors were penalized by deducting either 3 (drop out of field-of-view) or 1 (improper transfer) from the total number of objects moved.

Time-to-completion and errors committed were recorded for both pre- and post-tests. Data obtained from each 20-minute training session included number of full repetitions completed, number of individual objects moved, and errors committed. An overall score was calculated by subtracting the penalties accrued from errors from the total number of objects moved (Figure 2). Finally, “seconds-per-object” were calculated for the pre-test, six training sessions, and post-test in order to have all longitudinal assessments on the same measurement scale for analysis.

**Figure 2:**
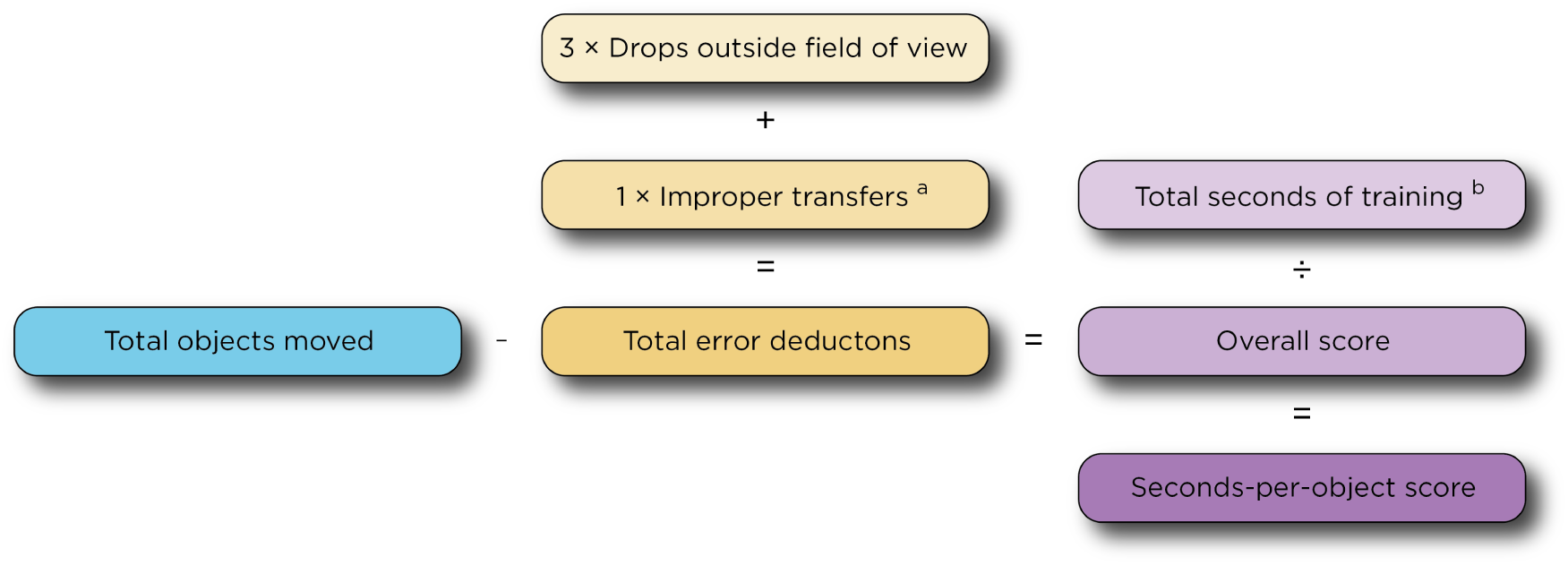
Scoring algorithm.

### F. Statistical Analysis

Baseline characteristics, personal behaviors, and prior surgical skill data were compiled and analyzed. Comparisons between the multiple intervention groups were performed with one-way analysis of variance (ANOVA). Comparisons between categorical variables were made with chi-square test for association. Individual performance on pre-tests and post-tests across the intervention cohorts was analyzed utilizing time-to-completion via independent samples *t*-tests and multiple one-way ANOVA tests.

Learning curves were plotted using the seconds-per-object scores for pre-test, six training sessions, and post-test. A second learning curve was produced including only the six training sessions to ensure any differences were not driven by differences on the pre-test between cohorts. These learning curves were normalized relative to the pre-test score and first training session, respectively. We analyzed the effects of intervention using linear mixed effects models. A repeated effect of training session was used to model the covariance of the residuals with a compound symmetry structure. In addition to the main effects of intervention and session, we also included the intervention-by-session interaction.

Sub-analyses applied independent samples *t*-tests to determine differences in performance metrics based on sex, handedness, and prior surgical experience. Finally, performance metrics were correlated to the various baseline characteristics and personal behaviors obtained on the questionnaire.

A *p*-value of < 0.05 was considered statistically significant. Statistical analysis was completed using R v3.5.0 (The R Foundation for Statistical Computing, Vienna, Austria).

## III. RESULTS

### A. Demographics

Sixty subjects (84.5% of screened participants) completed all six training sessions, including 20 each in the active bM1, active SMA, and sham (10 bM1, 10 SMA) tDCS groups (Figure 3). Among the three intervention groups, there was no difference in mean age (bM1 21.9 ± 5.2 vs SMA 23.5±5.4 vs sham 22.77±3.7 years), sex (males: bM1 8, SMA 4, sham 6), race (Caucasian: bM1 6, SMA 11, sham 8), and handedness (left: bM1 17, SMA 18, sham 18) (all *p* > 0.05).

**Figure 3:**
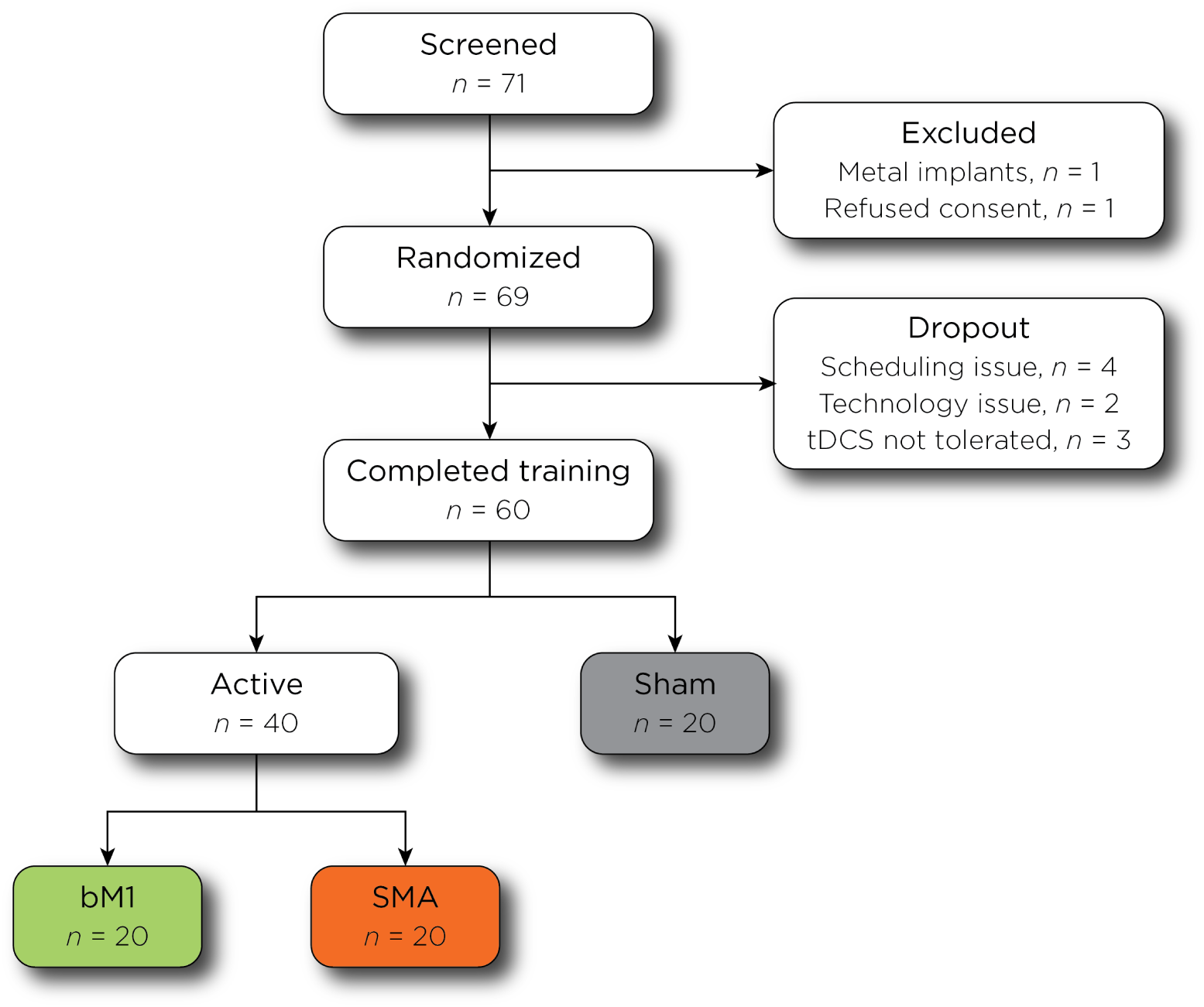
Consort diagram.

Six subjects had prior open surgical experience, and four had prior laparoscopic surgical experience. These individuals were distributed evenly amongst the three cohorts (open *p* = 0.574 and laparoscopic *p* = 0.765). Self-identified athletes (*n* = 34, 56.7%), musicians (*n* = 22, 36.7%), and artists (*n* = 10, 16.7%) were also equally represented across cohorts (all *p* > 0.05). However, video gamers had a larger presence within the bM1 cohort compared to SMA and sham (χ^2^ = 9.412, *p* = 0.009). There was no difference in the average amount of caffeine consumed (*p* = 0.15) or hours of sleep obtained (*p* = 0.24) prior to each training session across cohorts. Complete baseline characteristics are in Table 1.

**Table 1:** Baseline characteristics

|  | **Total**<br>(*N* = 60) | **bM1**<br>(*N* = 20) | **SMA**<br>(*N* = 20) | **Sham**<br>(*N* = 20) | ***p*-value<sup>†</sup>** |
| --- | --- | --- | --- | --- | --- |
| Age | 22.7 (4.8) | 21.9 (5.2) | 23.5 (5.4) | 22.7 (3.7) | *p* = 0.60 |
| Gender |  |  |  |  | *p* = 0.39 |
| Male | 18 (30%) | 8 (40%) | 4 (20%) | 6 (30%) |  |
| Female | 42 (70%) | 12 (60%) | 16 (80%) | 14 (70%) |  |
| Ethnicity |  |  |  |  | *p* = 0.86 |
| Hispanic | 7 (11.7%) | 2 (10%) | 3 (15%) | 2 (10%) |  |
| Non-Hispanic | 53 (88.3%) | 18 (90%) | 7 (85%) | 18 (90%) |  |
| Race |  |  |  |  | *p* = 0.70 |
| White | 25 (42%) | 6 (30%) | 11 (52%) | 8 (42%) |  |
| African American | 9 (15%) | 3 (15%) | 2 (10%) | 4 (21%) |  |
| Asian | 20 (33%) | 9 (45%) | 6 (29%) | 5 (26%) |  |
| Native American/Alaska Native | 0 (0%) | 0 (0%) | 0 (0%) | 0 (0%) |  |
| Native Hawaiian/Pacific Islander | 1 (1%) | 0 (0%) | 0 (0%) | 1 (5%) |  |
| More than one | 4 (7%) | 1 (5%) | 2 (10%) | 1 (5%) |  |
| Not reported | 1 (1%) | 1 (5%) | 0 (0%) | 0 (0%) |  |
| Handedness |  |  |  |  | *p* = 0.86 |
| Right | 53 (88.3%) | 17 (85%) | 18 (90%) | 18 (90%) |  |
| Left | 7 (11.7%) | 3 (15%) | 2 (10%) | 2 (10%) |  |
| Prior Open Surgery Experience | 6 (10%) | 1 (5%) | 2 (10%) | 3 (15%) | *p* = 0.57 |
| Prior Laparoscopic Surgery Experience | 4 (6.7%) | 1 (5%) | 2 (10%) | 1 (5%) | *p* = 0.77 |
| Self-Identification |  |  |  |  |  |
| Athlete | 34 (56.7%) | 11 (55%) | 12 (60%) | 11 (55%) | *p* = 0.94 |
| Musician | 22 (36.7%) | 5 (25%) | 9 (45%) | 8 (40%) | *p* = 0.39 |
| Gamer | 9 (15%) | 7 (35%) | 1 (5%) | 1 (5%) | *p* < 0.01 |
| Artist | 10 (16.7%) | 6 (30%) | 1 (5%) | 3 (15%) | *p* = 0.10 |
| Servings of Caffeine prior to Training |  |  |  |  | *p* = 0.15 |
| Day 1 | 0.45 (0.8) | 0.55 (0.8) | 0.2 (0.4) | 0.6 (1.1) |  |
| Day 2 | 0.45 (0.8) | 0.65 (0.9) | 0.2 (0.4) | 0.5 (0.9) |  |
| Day 3 | 0.45 (0.8) | 0.60 (1.1) | 0.25 (0.4) | 0.5 (0.7) |  |
| Hours of Sleep prior to Training |  |  |  |  | *p* = 0.24 |
| Day 1 | 6.8 (1.1) | 7.0 (1.2) | 6.6 (0.9) | 6.8 (1.1) |  |
| Day 2 | 6.9 (1.0) | 7.2 (1.1) | 6.7 (0.7) | 7.0 (1.0) |  |
| Day 3 | 6.8 (1.4) | 6.5 (1.4) | 6.6 (1.5) | 7.3 (1.2) |  |

All data reported as count (percent) or mean (SD)
<sup>†</sup>ANOVA for age, caffeine, and sleep; $\chi^2$ for remaining comparisons

### B. Raw Performance Metrics

All three cohorts had a gradual improvement in the average number of total repetitions completed (ranging from 10.3 to 18.4) and average number of total objects moved (ranging from 128.7 to 225.1) as they progressed through the 6 training sessions. Errors made by dropping the object outside of the field-of-view remained fairly stable throughout training, ranging from 0.7 to 1.8 across all cohorts. The average number of improper transfers gradually decreased in the SMA cohort (3.1 to 0.7) and had greater variability in the bM1 and sham cohorts. The overall scores accounting for objects moved and errors committed also increased over all six training sessions for all training cohorts (bM1 123.4 to 220.1, SMA 124.7 to 212.5, sham 130.5 to 208.5). Full, raw performance metrics can be found in Table 2.

**Table 2:**
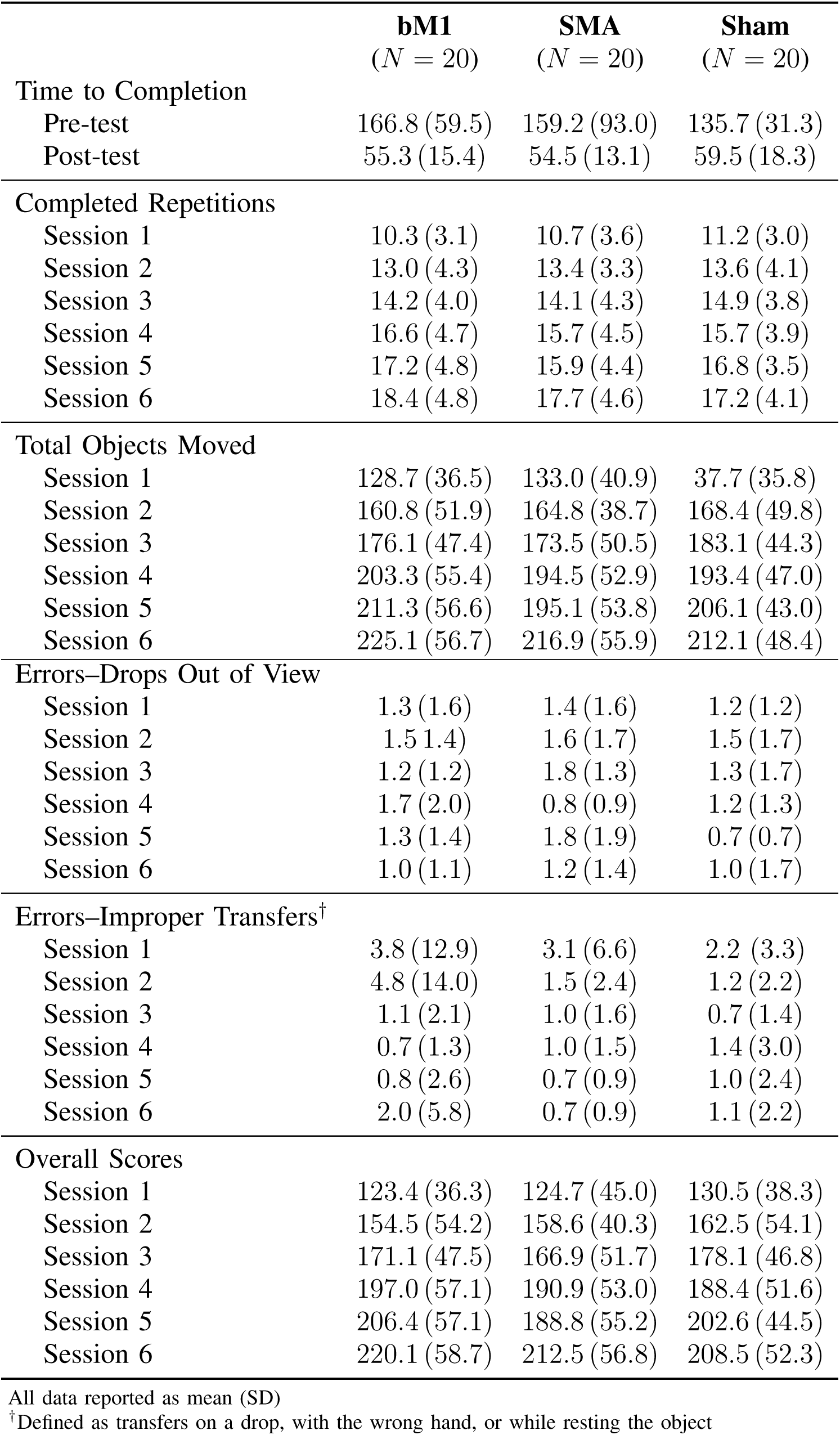
Average performance metrics by intervention group

|  | **bM1**<br>(*N* = 20) | **SMA**<br>(*N* = 20) | **Sham**<br>(*N* = 20) |
| --- | --- | --- | --- |
| **Time to Completion** |  |  |  |
| Pre-test | 166.8 (59.5) | 159.2 (93.0) | 135.7 (31.3) |
| Post-test | 55.3 (15.4) | 54.5 (13.1) | 59.5 (18.3) |
| **Completed Repetitions** |  |  |  |
| Session 1 | 10.3 (3.1) | 10.7 (3.6) | 11.2 (3.0) |
| Session 2 | 13.0 (4.3) | 13.4 (3.3) | 13.6 (4.1) |
| Session 3 | 14.2 (4.0) | 14.1 (4.3) | 14.9 (3.8) |
| Session 4 | 16.6 (4.7) | 15.7 (4.5) | 15.7 (3.9) |
| Session 5 | 17.2 (4.8) | 15.9 (4.4) | 16.8 (3.5) |
| Session 6 | 18.4 (4.8) | 17.7 (4.6) | 17.2 (4.1) |
| **Total Objects Moved** |  |  |  |
| Session 1 | 128.7 (36.5) | 133.0 (40.9) | 37.7 (35.8) |
| Session 2 | 160.8 (51.9) | 164.8 (38.7) | 168.4 (49.8) |
| Session 3 | 176.1 (47.4) | 173.5 (50.5) | 183.1 (44.3) |
| Session 4 | 203.3 (55.4) | 194.5 (52.9) | 193.4 (47.0) |
| Session 5 | 211.3 (56.6) | 195.1 (53.8) | 206.1 (43.0) |
| Session 6 | 225.1 (56.7) | 216.9 (55.9) | 212.1 (48.4) |
| **Errors–Drops Out of View** |  |  |  |
| Session 1 | 1.3 (1.6) | 1.4 (1.6) | 1.2 (1.2) |
| Session 2 | 1.5 (1.4) | 1.6 (1.7) | 1.5 (1.7) |
| Session 3 | 1.2 (1.2) | 1.8 (1.3) | 1.3 (1.7) |
| Session 4 | 1.7 (2.0) | 0.8 (0.9) | 1.2 (1.3) |
| Session 5 | 1.3 (1.4) | 1.8 (1.9) | 0.7 (0.7) |
| Session 6 | 1.0 (1.1) | 1.2 (1.4) | 1.0 (1.7) |
| **Errors–Improper Transfers<sup>†</sup>** |  |  |  |
| Session 1 | 3.8 (12.9) | 3.1 (6.6) | 2.2 (3.3) |
| Session 2 | 4.8 (14.0) | 1.5 (2.4) | 1.2 (2.2) |
| Session 3 | 1.1 (2.1) | 1.0 (1.6) | 0.7 (1.4) |
| Session 4 | 0.7 (1.3) | 1.0 (1.5) | 1.4 (3.0) |
| Session 5 | 0.8 (2.6) | 0.7 (0.9) | 1.0 (2.4) |
| Session 6 | 2.0 (5.8) | 0.7 (0.9) | 1.1 (2.2) |
| **Overall Scores** |  |  |  |
| Session 1 | 123.4 (36.3) | 124.7 (45.0) | 130.5 (38.3) |
| Session 2 | 154.5 (54.2) | 158.6 (40.3) | 162.5 (54.1) |
| Session 3 | 171.1 (47.5) | 166.9 (51.7) | 178.1 (46.8) |
| Session 4 | 197.0 (57.1) | 190.9 (53.0) | 188.4 (51.6) |
| Session 5 | 206.4 (57.1) | 188.8 (55.2) | 202.6 (44.5) |
| Session 6 | 220.1 (58.7) | 212.5 (56.8) | 208.5 (52.3) |

All data reported as mean (SD)
<sup>†</sup>Defined as transfers on a drop, with the wrong hand, or while resting the object

ANOVA calculated on the raw (non-penalized) completion times did not differ significantly between the bM1, SMA or sham cohorts at pre-test (166.8, 159.2, and 135.7 seconds respectively, *p* = 0.31) or post-test (55.3, 54.5, and 59.5 seconds respectively, *p* = 0.57). However, differences in improvement from pre-test to post-test completion times did differ significantly. The combined active tDCS cohort, bM1 and SMA, showed significantly greater improvement compared to sham (108.1 vs 76.3 seconds, *p* = 0.018). Similarly, the bM1 active (111.5 vs 76.3 seconds, *p* = 0.022) and the SMA active (104.7 vs 76.3 seconds, *p* = 0.015) conditions each, alone, showed significantly greater improvement relative to sham (Figure 4).

**Figure 4:**
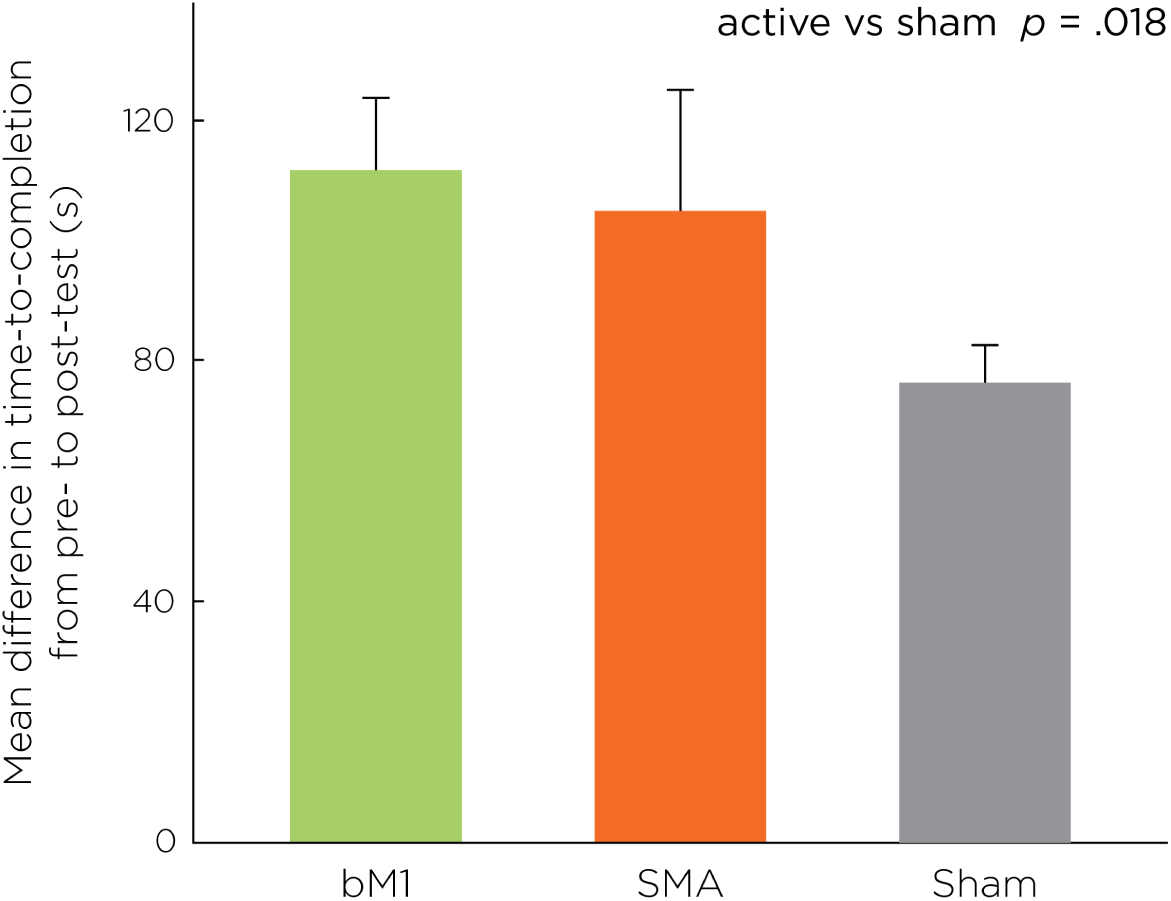
Mean improvement in time-to-completion (seconds) from pre-test to post-test for each of the three intervention groups. Whiskers denote standard error of the mean. Significant pairwise differences are present between each active group and the sham group.

### C. Learning Curves

Linear mixed effects modeling on the scored performance data (total objects moved minus errors) demonstrated that all subjects, regardless of intervention group, demonstrated learning across the six training sessions (Figure 5). A significant group by session interaction was observed. There was no significant difference in performance when comparing the bM1 and SMA cohorts (*t* = 0.043, *p* = 0.97). This difference between active and sham tDCS remained when looking solely at the six 20-minute training sessions, with a statistically significant difference between bM1 and sham cohorts (*t* = 2.07, *p* = 0.039), but not between the SMA and sham cohorts (*t* = 0.85, *p* = 0.40).

**Figure 5:**
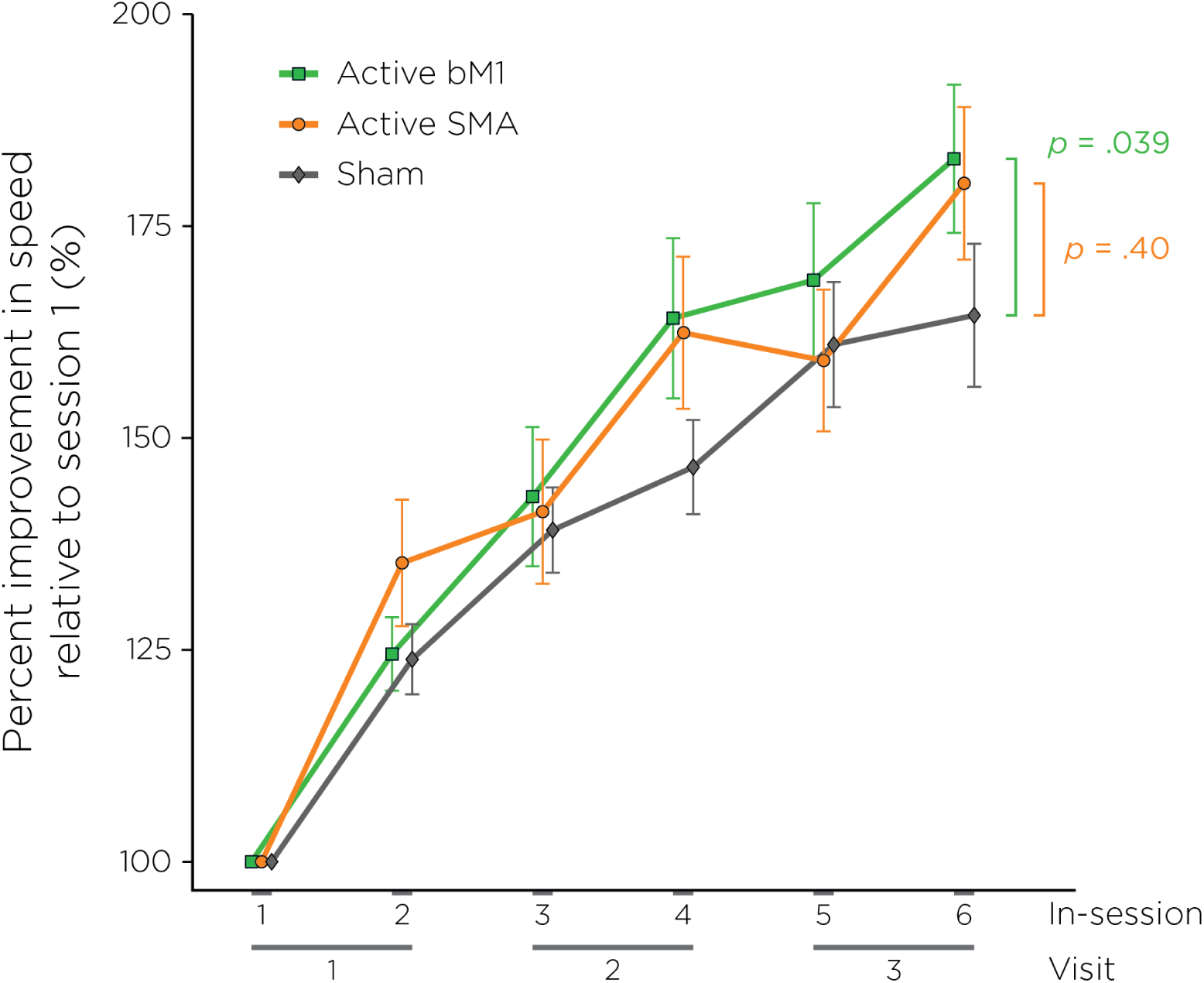
Learning curves stratified by intervention group and reported as percent improvement in seconds-per-object normalized to the session 1 performance. Whiskers denote standard error of the mean.

### D. Performance Variability

Comparing performance between male versus female subjects and right versus left handed subjects did not reveal significant differences in pre-test, post-test, improvement from pre- to post-test, or overall score during the six training sessions (all *p* > 0.05). Similarly, no differences were observed in performance between individuals who self-reported as athletes, musicians, artists, or video gamers, versus those reporting not to belong to these classifications (all *p* > 0.05). Age, servings of caffeine, and hours of sleep did not correlate with any of the task metrics (all *p* > 0.05).

Similarly, subjects with prior open surgery experience had no difference in performance compared to those without prior experience (*p* > 0.05). However, subjects with prior laparoscopic surgery experience had significantly better performance in the second (*t* = 4.1, *p* < 0.001) and third (*t* = 3.7, *p* = 0.001) 20-minute training sessions compared to subjects without laparoscopic surgery experience.

### E. Compliance and Adverse Events

Eleven participants (15.5%) who were screened did not complete the entire study protocol. One participant did not consent while another was excluded at screening due to metal implants. Six participants withdrew due to technology issues and scheduling conflicts. Three participants had intolerance of tDCS due to lightheadedness leading to early cessation of training and exclusion from analysis. Two of these three subjects were in an active tDCS cohort; one subject revealed she had nothing to eat throughout the day prior to the study session while the other subject disclosed she ran 10 miles directly prior to the study session. The third intolerant subject was in the sham cohort and experienced dizziness during the initial ramp up and final ramp down during the second training session.

## IV. DISCUSSION

We present the largest to date, pre-registered, randomized, and sham-controlled study of the effectiveness of tDCS coupled with simultaneous surgical skills training, and the first study to contrast two different tDCS electrode configurations (bilateral primary motor cortex or the supplementary motor area). Training coupled with active tDCS exhibited significantly greater learning in the Fundaments of Laparoscopic Surgery Peg Transfer Task compared to sham tDCS for both electrode configurations although the SMA configuration produced greater variability. The results from this pilot study demonstrate the potential of accelerating laparoscopic technical skill acquisition through the addition of tDCS. Given the safety and ease of administering tDCS along with these promising preliminary results, this novel technique could be a useful and practical augmentation to current technical skill training within surgical residency.

All general surgery residents utilize deliberate practice within the surgical simulation lab in order to pass the national, standardized FLS curriculum allowing for board eligibility. However, the amount of time allotted for training is restricted by the maximum 80-hour work week. Our results suggest trainees could achieve faster and greater learning with the application of tDCS during practice sessions. Subjects training with active tDCS reached the same level of technical skill in roughly two thirds the time as compared with sham tDCS. This translates to about 40 minutes less total training time when using active tDCS, a savings that could likely be compounded when factoring in training on all five FLS tasks. Collectively, the active tDCS cohorts improved an average of 31 seconds more per repetition of the Peg Transfer Task compared to the sham cohort. This is a highly meaningful amount considering a total completion time of 48 seconds is considered expert proficiency in this particular task [37].

There is only one prior study utilizing tDCS within the field of surgery [32]. In that study, Ciechanski and colleagues applied unilateral, anodal tDCS within neurosurgery skills training using a simulated tumor resection model. They concluded tDCS safely enhanced simulation-based neurosurgical skill acquisition with larger gains appreciated in the lower skill participants. A strength of their study included a retention test at 6 weeks post-training which revealed no skill decay for either training group. Unfortunately, the tDCS intervention was fairly limited with only a 24-minute training session consisting of 8 task repetitions. The current study applying tDCS within general surgery skill training utilizes a reliable and validated model in the FLS Peg Transfer Task, with substantially more accrued training time. Our methodology included a training curriculum designed to prevent an early plateau in performance and allow enough training over time for the development of sufficient learning curves to determine the effect of our intervention. These studies together support the rationale to continue evaluation of tDCS as an augmentation to technical skill training within the surgical simulation lab.

Other strengths of our study include sham-control, comparison of two electrode configurations, E-field simulation, and use of a device permitting complete double-blinding of subjects and investigators to active versus sham conditions. A recent tDCS consensus manuscript by Buch and colleagues [26], proposed a checklist with standards for tDCS studies to allow for methodological reproducibility. The current study was designed prior to the publication of this updated list, but it fulfills all but a few of the suggested criteria.

Limitations of this study include the potential that pre-test differences in performance may have influenced the results. While not statistically significant, the sham groups had better performance at baseline, which may have contributed to less change in performance due to ceiling effects. A baseline difference was not anticipated given the overall sample size of 60 individuals and random group assignments. Nonetheless, future studies may wish to adopt a stratified randomization approach to mitigate unexpected baseline difference.

Of note, the learning curve of the SMA cohort had more variability compared to the bM1 cohort. This likely contributed to the lack of significant differences when considering learning in the absence of pre-test and post-test performance. The SMA electrode configuration includes cathodal placement over the frontal cortex which could theoretically impair subjects’ attention or cognitive control [38]. This possibility, and the optimal electrode configuration, current strength, duration of stimulation, and timing of stimulation relative to manual-skill training merits further investigation.

This was a pilot study with no prior investigation to allow for a power calculation to determine sample size. Therefore, 20 subjects were randomized into each cohort in concordance with prior tDCS research studies [26]. The current study included both right handed (*n* = 53) and left handed (*n* = 7) subjects, but the anode and cathode electrodes were placed in the same position regardless of handedness. Given the bimanual nature of the task future studies may wish to quantify the lateralized effects of anodal and cathodal stimulation or use unipolar motor system montages. Finally, the current study utilized a single simulation task leaving the generalizability of this intervention yet to be determined. However, tDCS did provide a training advantage, and dexterity utilized in the Peg Transfer Task also applies to other surgical skills including open, endoscopic, and robotic tasks.

Moving forward, future studies will be needed to replicate these findings, and determine if similar benefits can be gained with surgical residents who have already achieved a higher baseline level of surgical knowledge and skill compared to the studied cohort. In addition, translation of these findings to a commercially available tDCS device would also allow more feasible implementation into the surgical skills laboratory, allowing training without research supervision. While outside the scope of the current study, tDCS could be useful to accelerate remediation programs for trainees found to be outliers in regards to their technical skills.

## V. CONCLUSION

The results of this study suggest the acquisition of basic laparoscopic skill can be enhanced with the addition of active tDCS during training. This finding has the potential to accelerate general surgery training within the current climate of work hour restrictions and limited time for the development of technical skills within competency-based curricula.

## ACKNOWLEDGMENTS & DISCLOSURES

This study was presented as an oral presentation at the Innovative Techniques session at the 2018 Annual ACS Surgical Simulation Summit: An International Multi-Professional Meeting in Chicago, IL on March 16–17, 2018.

This study was supported internally at our institution. M. L. Cox is supported by a National Institutes of Health T32 Training Grant with grant number T32HL069749. Z.-D. Deng is support by the National Institute of Mental Health Intramural Research Program and a NARSAD Young Investigator Award from the Brain & Behavior Research Foundation.

The authors do not have any commercial, proprietary or financial interest in any device, equipment, instrument, or drug related to this article.

S. H. Lisanby contributed to this article while at Duke University, prior to joining NIMH. The views expressed are her own and do not necessarily represent the views of the NIH or the United States Government.

